# Non-triplet genetic code in *Euplotes* ciliates is a result of neutral evolution

**DOI:** 10.1101/2022.10.12.511967

**Authors:** Sofya Gaydukova, Mikhail Moldovan, Adriana Vallesi, Stephen M. Heaphy, John F Atkins, Mikhail S. Gelfand, Pavel V. Baranov

**Affiliations:** Skolkovo Institute of Science and Technology, Moscow, 121205, Russia; Faculty of Bioengineering and Bioinformatics, Lomonosov Moscow State University, Moscow, 199911, Russia; Laboratory of Eukaryotic Microbiology and Animal Biology, School of Biosciences and Veterinary Medicine, University of Camerino, Camerino, 62032, Italy; School of Biochemistry and Cell Biology, University College Cork, Cork, T12 XF62 Ireland; Department of Human Genetics, University of Utah, Salt Lake City, UT 84112; A. A. Kharkevich Institute for Information Transmission Problems RAS, Moscow, 127051, Russia

## Abstract

Although several variants of the standard genetic code are known, its triplet character is universal with an exception in ciliates *Euplotes*, where stop codons at internal mRNA positions specify ribosomal frameshifting. How did *Euplotes* spp. evolved and maintained such an unusual genetic code remains a mystery. To investigate these questions, we explored the evolution of frameshifting occurrence in Euplotes genes. We sequenced and analyzed several transcriptomes from different *Euplotes* spp to characterize the gain-and-loss dynamics of frameshift sites. Surprisingly, we found a sharp asymmetry between frameshift gain and frameshift loss events with the former exceeding the latter by about 10 folds. Further analysis of mutation rates in protein-coding and non-coding regions revealed that this asymmetry is expected based on single nucleotide mutation rates and does not require positive selection for frameshifting. We found that the number of frameshift sites in *Euplotes* spp is increasing and is far from the steady state. The steady equilibrium state is expected in about 0.1 to 1 billion years leading to about a 10 fold increase in the number of frameshift sites in Euplotes genes.

---

The sequential non-overlapping triplet nature of genetic decoding was established by Crick, Brenner and their colleagues in the early 60s^1^. Almost all proteins are encoded by such sequential nucleotide triplets, codons. Thus, the decoding ribosome moves along mRNA in one of the three-periodic phases known as the reading frames. Errors in maintaining the reading frame are more detrimental than missense errors as they affect the entire downstream sequence of the protein^2^. Consequently, spontaneous shifts between reading frames are highly infrequent^3^. As the accuracy of triplet decoding is sequence-dependent, frameshifting-prone sequences are selected against in protein-coding genes^4,5^. However, the sequence dependence of frameshifting efficiency enabled evolution of genes that exploit this phenomenon to regulate their expression in the process known as programmed ribosomal frameshifting^6^. Although genes requiring ribosomal frameshifting for their expression have been found in most organisms, such genes are usually extremely rare, though common in viruses^7^ and transposable elements^8^. To achieve higher efficiency, programmed ribosomal frameshifting often requires the presence of elaborate stimulatory signals such as RNA pseudoknots altering progression of the ribosome^9,10^ or nascent peptides interfering with the ribosome function from within^11,12^. Even with the assistance of such stimulators, the efficiency of ribosomal frameshifting is usually lower than that of the competing triplet decoding^6^. The product of ribosomal frameshifting is synthesized in addition to the product of standard translation.

Ribosomal frameshifting observed during mRNA translation in ciliates of the genus *Euplotes*^13–17^ is in a striking contrast with programmed ribosomal frameshifting. It has been proposed to occur whenever a stop codon is encountered by the ribosome (Figure 1a). Unlike programmed ribosomal frameshifting it is highly efficient with effectively no products of termination being detected at the stop codons at internal positions^13^. Termination of translation occurs only near the 3’ ends of mRNAs in a close proximity to the polyA tails^13^ similarly to the situation in the species where all three stop codons have been reassigned to code for amino acids^18–20^ as outlined in ref. ^21^. Therefore, ribosomal frameshifting has been suggested to be a part of the standard Euplotes genetic code^13^ (Figure 1c).

**FIGURE 1.**
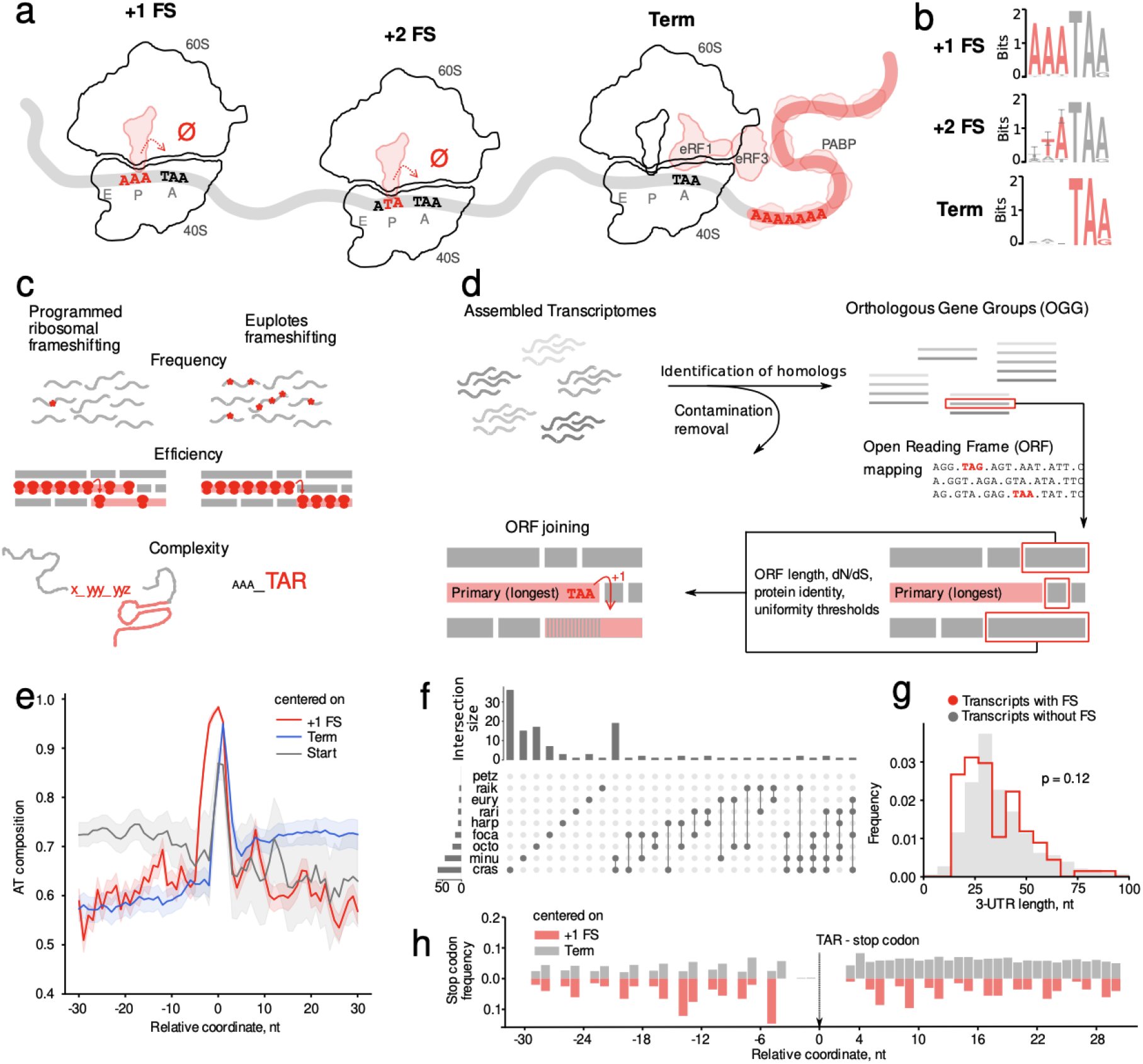
Euplotes transcriptomes and identification of frameshift sites. **(a)** The proposed mechanism of ribosomal frameshifting and translation termination in *Euplotes* spp. **(b)** Sequence Logos of frameshift sites and terminating stop codons contexts. **(c)** Properties of ribosomal frameshifting in *Euplotes* spp. in contrast to programmed ribosomal frameshifting. **(d)** Schematic of the algorithm for the identification of frameshift sites. **(e)** AT composition relative to start and stop codons (FS and terminators) centered at zero. **(f)** UpSetR representation of the distribution of identified frameshift sites across species. Rows correspond to the species and columns to the number of unique or shared frameshift sites. **(g)** Distributions of 3’ UTR lengths (nt) in transcripts with (red) and without (grey) frameshifts. **(h)** Stop-codon frequency around frameshifting (red) and terminating (grey) stop-codons.

It is not clear, however, how such a non-triplet feature of the *Euplotes* genetic code has evolved and which processes enable its persistence. To address these intriguing questions, we explored the evolution of frameshift site (FS) occurrences across a group of *Euplotes* sequences.

## Detection of frameshift sites

We have sequenced transcriptomes of nine species from six major clades of the *Euplotes* phylogenetic tree (Extended Data Figure 1): freshwater *E. octocarinatus*, brackish waters *E. harpa*, and marine *E. focardii, E. petzi, E. euryhalinus, E. rariseta, E. minuta, E. raikovi*, and *E. crassus*. The transcriptomes were assembled and combined into 1614 orthologous groups containing 4903 sequences (Methods). In all subsequent analyses, we considered only transcripts for which orthologs could be identified as we would not be able to assess the evolutionary trajectories of FSs from individual sequences. This also reduces contamination with sequences derived from other organisms found in the environment (Methods).

To detect FSs in these transcriptomes we developed a systematic and unbiased procedure. We assumed absolute specificity of frameshifting in *Euplotes* spp., which means that ORFs following a frameshifting event are translated in a single reading frame. The procedure starts with the longest ORF that is then extended with adjacent or overlapping ORFs joined with either +1 or +2 frameshifting or stop codon readthrough (Figure 1d). Candidate ORFs were detected using stringent criteria relying on minimal ORF length, high sequence similarity with their orthologs, signatures of purifying selection typical for the evolution of protein-coding sequences, and the uniformity of the two latter characteristics along the candidate protein-coding sequence. To avoid arbitrariness in thresholds underlying these criteria, we considered these thresholds as parameters and fine-tuned them to obtain robust results. Subsequent validation using ribosome profiling data generated in *E. crassus*^4^ demonstrated high specificity of the algorithm, with true- and false-positive discovery rates of 54% and 0%, respectively. Using this approach, we identified translated regions in the generated transcriptomes and found 3.9% of transcripts to contain FSs (8.3% orthogroups), i.e. 197 instances of +1 and 16 instances of +2 frameshifting sites distributed across 192 sequences from the total 4903 (Figure 1bf). No stop codon readthrough events could be identified. No frameshifting events were also identified in *E. petzi*, though manual analysis of alignments allows to identify a few events of frameshifting. Thus we suggest that this result is due to overly stringent criteria and a small sample of *E. petzi* transcripts that had an ortholog compared with other species. The set of predicted FSs and the predicted terminating stop codons (terminators) confer the previously reported features of *Euplotes* coding sequences, such as the prevalence of +1 shifts (predominantly at AAA preceding stop codons), short 3’ UTRs, enhanced AT-content in 3’ and 5’ UTRs and high frequency of stop codons in 3’ UTRs (Figure 1egh).

## Evolution of frameshifting in *Euplotes* spp

The unusually large number of tolerated, highly efficient FSs in *Euplotes* spp, a feature not observed in any other group of cellular organisms studied so far, calls for an evolutionary explanation. Although the current amount of data renders any population-based methods inefficient in this case, we can assess some basic evolutionary features of FSs, such as their general effects on fitness, from the FS gain and loss rates along the phylogeny.

We inferred frameshift site gain and loss events based on the reconstructed ancestry (Figure 2a). We found that the frequency of gains exceeds that of losses by about ten fold (39 vs 4). This sharp asymmetry may be explained by positive selection, however, to test for selection we have to consider probabilities of FS gains and losses rather than the numbers of respective events. These probabilities depend on the number of contexts suitable for FS gains and the number of existing FSs (that may be lost). The evolution of FSs is likely to be context-dependent since ~72% of all gain and loss events occurred due to the insertion or deletion of T in AAA_[T]AR (underscore separates codons, R=A/G). Thus, we initially focused on the analysis of the evolution of FSs conforming to this specific pattern. We define the probabilities of frameshift gains and losses as *P_g_* = *n_i_/K* and *P_l_* = *n_d_/F*, respectively, where *n_i_* is the number of observed insertions of T yielding new FSs, *K* is the number of ancestral AAA_AR sequence motifs, *n_d_* is the number of observed deletions of T leading to the loss of FSs, and *F* is the number of AAA_TAR FSs. As opposed to simple counts of FS gains and losses, the FS loss probability exceeds the FS gain probability by 3 to-30-fold (Figure 2d, *p*=0.028, the permutation test). This discrepancy may be explained by selection against novel FSs combined with increased mutational pressure favoring FS gains. Concerning the latter, we observe the numbers of FS gain contexts to be on average 74-fold higher than the numbers of existing FSs. This inflates the probability of FS-gain mutations compared to FS losses^22^. The selection against FSs may be estimated from the differences in the observed probabilities of FS gain and loss (Methods), with the selection-to-drift balance calculated as the average *S = 4sN_e_* of FSs, where *s* is the selection coefficient and *N_e_*, the effective population size.^22^

**FIGURE 2.**
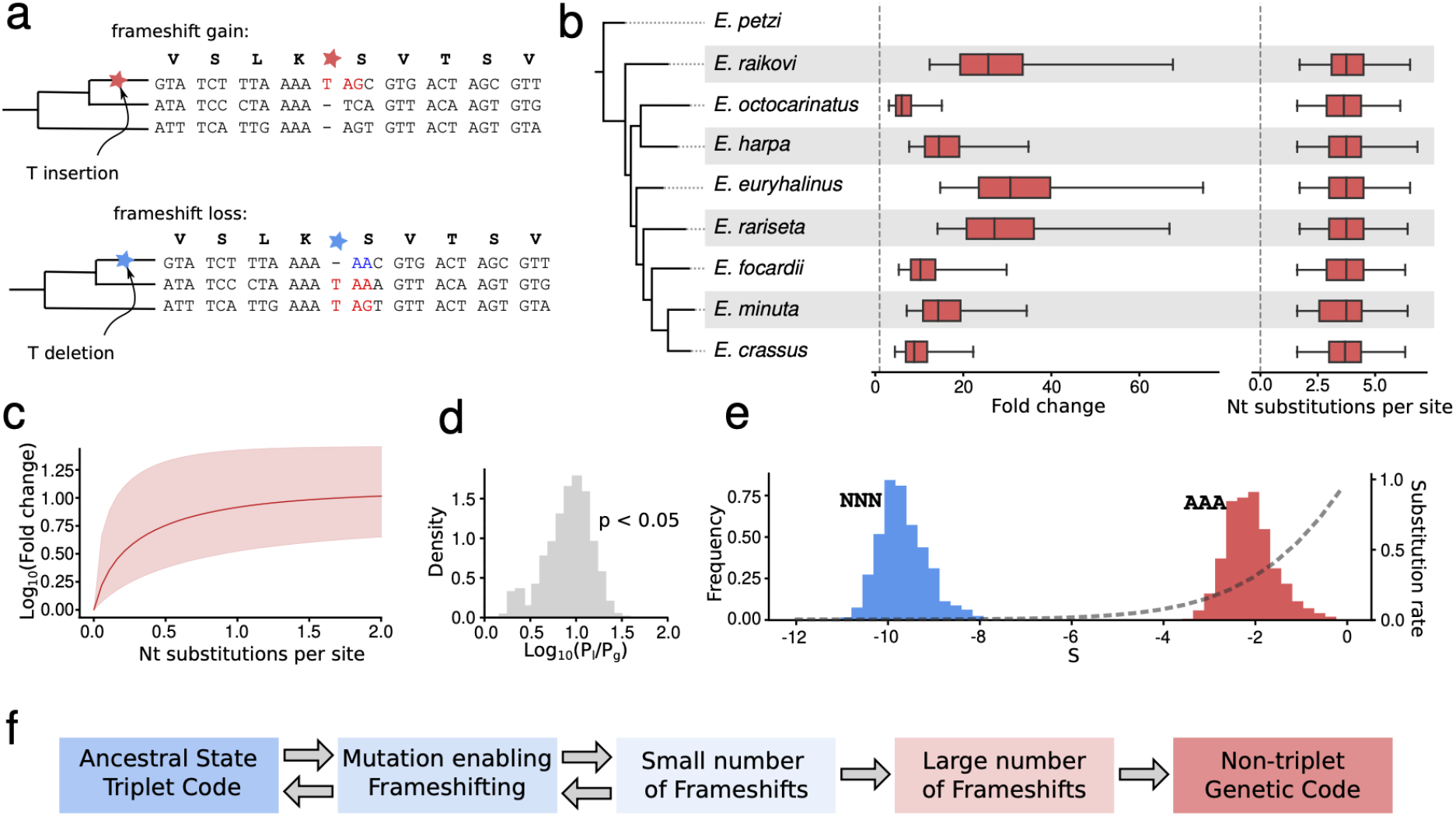
Increasing numbers of frameshift sites during the evolution of *Euplotes* spp. **(a)** Inference of gains (top) and losses (bottom) of FSs based on the ancestral states. **(b)** (left) Phylogenetic tree of studied *Euplotes* species. (middle) Distributions of the fold changes in FSs numbers upon reaching equilibrium. (right) Distributions of time intervals (in nucleotide substitutions) required to reach 95% of the FSs number expected at the equilibrium. **(c)** Projected fold change of the number of frameshift sites over time. Red shading indicates the 95% confidence interval obtained from permutations. **(d)** Ratio of FS gain and loss probabilities for AAA_[T]AR contexts. **(e)** Permutation distributions of *S* values for selection against FS arising in AAA_[T]AR contexts (red) and of upper bounds for *S* in non-AAA_[T]AR contexts (blue). Dashed line is the theoretical dependence of the normalized substitution rate for variants affected by selection and genetic drift on the *S* value. **(f)** Proposed scheme of the emergence and subsequent entrenchment of the non-triplet genetic code.

The calculated *S* (Figure 2e) differed for frameshifts in the AAA_[T]AR and NNN_[T]AR contexts, and while *S* for frameshifts in the latter context are in the highly deleterious range (upper bound *S*=-10±1, the confidence interval derived from permutation analysis), frameshifts in AAA_[T]AR context seem to be only slightly deleterious (S=-2±1), which is consistent with fitness-reducing FS effect in *Euplotes* spp. that may be due to ribosome pausing^13^. Indeed, the observed *S*=-2 may result in both negative selection and drift influencing accumulation of mutations, whereas *S*=-10 indicates efficient negative selection against FSs arising in NNN_[T]AR contexts^22^.

Thus, the large numbers of contexts suitable for FS gains compared to the numbers of already existing FSs yield an increased mutational pressure towards FS accumulation, which counters the weak selection against FSs in favorable contexts. But are these processes at equilibrium and, consequently, are the frequencies of euplotid FSs constant in time? And, if there is no equilibrium, how distant are *Euplotes* spp. from it? The following differential equation describes the change of the number of FSs over time where *u_g_* and *u_l_* are, respectively, the rates of gain and loss derived from their probabilities (see Methods), and *K* and *F* are, as earlier, the (constant) number of ancestral AAA_AR sequence motifs and the (variable) number of current AAA_TAR FSs, respectively:

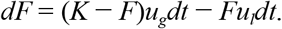

By solving this equation (see Methods) we projected the changes in the number of FSs over time. At infinite time, this function reaches the upper asymptote which corresponds to the number of FSs when the gains and losses are at equilibrium. We find the asymptote to be 5-25-fold (depending on the species) larger than the current numbers of FSs (*p*<0.0001, the permutation test, see Methods) (Figure 2b, red).

Next, we estimated the time needed for reaching the equilibrium. As formally this time is infinite, we consider the effective equilibrium as the time point when the number of FSs is 95% of its asymptotic value. We found these times to be consistent across species and constitute 1.7-6.3 nucleotide substitutions per site on average (Figure 2b), which is about 0.68-2.52-fold larger than the age of the considered group of *Euplotes* spp.^23^. Thus, our results indicate that the euplotid FSs are only in an early stage of the accumulation process.

The number of euplotid FSs per genome at the equilibrium is expected to be between 17131 and 71685. Thus, if we consider independent effects of FSs on fitness, the net lag load (loss of relative fitness) *L* conveyed by the total body of FSs becomes *L = 1 - (1-S/N_ρ_)^F^*, which, depending on the *N_e_* value, may be either substantial and pose a potential hazard for the survival of *Euplotes*, or negligible. To assess this,, we calculated the dependence between the net lag load of eventual frameshifts calculated from the obtained per-site 4*sN_e_* values and *N_e_*. We observe that the net lag load would drastically decrease fitness of *Euplotes* spp. on *N_e_*<10^5^, whereas on *N_e_*>10^6^ there is only a minor fitness decrease (Extended Data Figure 4). Although the exact *N_e_* values of *Euplotes* spp. cannot be calculated in the absence of micronuclei genomic data, we may assume the euplotid *N_e_* values to be larger than 10^6^, as smaller organisms typically have much larger *N_e_* (for comparison, *N_e_* ~ 10^6^ for the current human population^24^). Thus, although there will eventually be some significant load associated with frameshifts, it should not impact the survival of *Euplotes* spp.

## Conclusions

The findings presented here suggest that *Euplotes* spp. are only in the beginning of their evolution towards a balanced non-triplet genetic code. Since the ability of efficient frameshifting at internal stop codons is shared among all *Euplotes* spp., it is likely to have emerged in their common ancestor. However, as we showed here, the process of FS accumulation is very slow and the number of FSs observed at present time constitutes only about 4-20% of its expected maximum at the equilibrium.

Is frameshifting itself useful for *Euplotes* spp.? Does it increase their fitness and at least partially compensate for the detrimental effects of FSs? It has been suggested that alternative genetic codes are frequently used in ciliates to protect their genomes from foreign genetic elements^26^. In addition, efficient frameshifting at out-of-frame stops in *Euplotes* spp. makes their genes resistant to single nucleotide insertions in the protein-coding regions. Indeed, we found several instances of insertions that disrupt the protein coding reading frame, but then the reading frame is restored due to frameshifting at a premature stop codon downstream (Extended Data Figure 5). Thus, single nucleotide insertions in *Euplotes* coding sequences result in a change of only a short segment of the encoded protein (between insertion and newly formed premature stop codon) rather than the entire protein sequence encoded downstream.

In addition to these and other potential possibilities of frameshifting “benefits”, our study suggests an alternative more likely evolutionary explanation of the abundant frameshifting in *Euplotes*. Once a species develops an ability to frameshift ribosomes at in-frame stop codons with high efficiency, the number of in-frame stop codons would start growing even if frameshifting is mildly deleterious. Once a certain number of FSs is reached, the process is expected to become entrenched and hence irreversible, as a reversal of the change that enabled frameshifting would result in mistranslation of many genes (Figure 2f). Hence, the high number of FSs in *Euplotes* does not imply that they are beneficial, but simply that they are not a limiting factor in evolution and are not, individually or collectively, under strong selection. Thus we conclude that changes in the genetic code, even as profound as the violation of its triplet character, may be the result of neutral evolution.

## METHODS

### Species selection and cell cultures

*Euplotes* strains used in this study are part of a vast collection maintained in the laboratories of the Universities of Pisa and Camerino. Each species was selected on the basis of the position it occupies within the *Euplotes* phylogenetic tree, which is commonly regarded as articulated in six major clades (numbered I to VI from the bottom) (Extended Data Figure 1)^25,26^. *E. petzi* forms the most basal clade I. *E. raikovi* clusters with few other species into clade IV. *E. octocarinatus* and *E. harpa* lie in the well-supported clade V which included most of the freshwater *Euplotes* species, but they split in two different subclades. *E. euryhalinus, E. rariseta, E. minuta, E. crassus* and *E. focardii* cluster together into the poorly resolved and species-richest clade VI. However, these species branch into three different subclades. One subclade includes *E. euryhalinus* and *E. rariseta;* a second subclade includes *E. focardii* and a third subclade includes *E. minuta* and *E. crassus* which, among the all the species of this study, are the most closely related as also certified by previous breeding investigations^27^.

The selected *Euplotes* species have different ecologies. *E. octocarinatus* and *E. harpa* are temperate freshwater and brackish species, respectively^28,29^, while all the others are marine species. *E. focardii* is endemic to Antarctica^30^. *E. petzi* and *E. euryhalinus* have a bi-polar (Antarctic and Arctic) distribution^31,32^. *E. rariseta, E. crassus, E. minuta* and *E. raikovi* are virtually ubiquitous in temperate coastal areas^33^.

### RNA preparation and sequencing

Cultures were grown under a daily cycle of 12 h of dark and 12 h of very weak light, at 4–6 °C (polar species) or 18–20 °C (non-polar species), using as food the green algae *Dunaliella* (marine species) and *Chlorogonium* (freshwater species). They were expanded by daily food additions up to a cell density of about 10^4^cells/ml, then washed free of food and debris, and re-suspended for 3 (temperate species) or 6 (polar species) days in fresh marine or distilled water before being harvested. The TRIzol plus purification kit (Thermo Fisher Scientific) was used to purify total RNA, following the manufacturer’s recommendations. Samples of about 10^7^ cells were concentrated by mild centrifugation and lysed by rapid resuspension in 1 ml TRIzol reagent containing phenol and guanidine. After chloroform addition and centrifugation, an equal volume of 70% ethanol was added to the aqueous phase containing RNA, which was next purified using silica cartridges. As a rule, an on-column DNAse treatment was carried out for 45 min at room temperature to obtain DNA-free RNA preparations. After washing, RNA was eluted with 30 μl RNAse-free water and stored at −80 °C before use. RNA concentration and purity were estimated by NanoDrop One Spectrophotometer (Thermo Fisher Scientific), while RNA integrity was analyzed by agarose gel electrophoresis.

cDNA library preparation and sequencing was carried out by BGI using Illumina TruSeq library construction and sequencing at Illumina HiSeq 2000 using 101PE.

### Assembly of transcriptomes and construction of orthologous gene groups (OGG)

The resulting read libraries were trimmed with Trimmomatic^34^ with parameters: ILLUMINACLIP:TruSeq3-SE:2:30:10 LEADING:3 TRAILING:3 SLIDINGWINDOW:4:15 MINLEN:36. The transcriptomes were assembled with Trinity^35^ using the default set of parameters.

Orthologous gene groups (OGGs) were constructed using ProteinOrtho v. 5.15^37^ with parameters: -p=blastn -e=1e-25 -identity=70 on obtained transcriptomes.

The constructed OGGs were aligned with MUSCLE^38^ with default parameters. For the coding region alignment, we employed TranslatorX^39^ with parameters: -p M -t F -w 1 -c 10. Protein alignments of predicted proteins were constructed with the Smith-Waterman algorithm^40^.

To determine whether a sequence of a transcript is in sense or antisense orientation we relied on the locations of polyA and polyT tails, polyA tails at the 3’ ends were used as indicator of the sense strand, while transcripts with polyT were classified as antisense and reverse complements of these transcripts were used instead. In the absence of polyA/T tails (truncated transcripts), we choose the orientation yielding the lowest number of indels in the longest ORF aligned to one of its orthologs.

### Calculation of identity and *d_N_/d_S_* values

To calculate the protein identity between two ORFs, they were translated using the euplotid genetic code (#10^41^) and aligned with the Smith-Waterman algorithm^40^. All gap-containing positions were removed. The identity was calculated as the number of identical amino acids divided by the number of non-gap positions.

We employed two ways to calculate the *d_N_/d_S_* ratio: the Nei-Gojobori method^42^, which yields the values which can be straightforwardly compared by simple statistics (see below) and the Nielsen-Yang method^43^ for the correction and validation of the obtained *d_N_/d_S_* values as implemented in PAML^44^.

For the analysis of *d_N_/d_S_* uniformity was performed from compared ORFs^45^. At each permutation round all four values needed to calculate the *d_N_/d_S_* ratio, that is *dN, N, dS, S* (in the Nei-Gojobori notation^42^), were sampled from the respective Poisson distributions with the parameters derived from the data. The *d_N_/d_S_* ratios obtained for two ORFs were then compared. The percent of permutations which resulted in *d_N_/d_S_* of adjacent (shorter) ORF being higher than *d_N_/d_S_* of the primary (longer) ORF was used as the metric of uniformity.

### Transcriptome quality assessment

Ciliates are obligate heterotrophs^46^ mostly feeding on algae and other microorganisms, the remnants of which remain in the ciliate cytoplasm. They also may have intracellular symbionts^47^. Hence the experimental separation of the ciliate DNA from the DNA of symbionts and prey is currently not feasible^48–50^. This necessitates removal of contaminant sequences from assembled transcriptomes. We employed a four-step filtering procedure with subsequent controls to ensure that the transcripts considered in downstream analyses were indeed ciliate transcripts:

1. AT-content was calculated for every transcript and the resulting distribution was assessed (Extended Data Figure 3). Since some distributions were bimodal, we filtered out all transcripts with the AT content less than 4 standard deviations from the distribution mean. This constraint follows observations that ciliate transcriptomes are AT-rich^51^.
2. All transcripts having no homologs in other transcriptomes obtained in this study (singletons) were filtered out (see “Assembly of transcriptomes and construction of orthologous gene groups” above).
3. For each sequence, the nucleotide identity with its closest homolog from the sample was calculated. The resulting distribution appeared sharply trimodal (Extended Data Figure 4) with ciliate sequences found only in the middle peak, as verified with BLASTn search (see below). From this distribution, we obtained the boundaries of the middle peak corresponding to the between-ciliate identity range. Then we discarded all transcripts with identity below the obtained lower bound of 0.65 and above the obtained upper bound of 0.95, which corresponded to the boundaries of the middle peak.
4. For the remaining sequences the taxonomy of candidate contaminating species was identified. For this, we randomly selected 1000 transcripts from each transcriptome and performed online BLASTn^52^ search against Genbank non-redundant database using the NCBI implementation at https://blast.ncbi.nlm.nih.gov/Blast.cgi. A non-ciliate species was considered as being closely related to a contaminating species if it produced the best hit with at least 50% identity and alignment length of at least 50% relative to the query at E-value below 10^-5^. A total of 41 contaminating (or closely related) organisms were identified (Extended Data Table 2). All transcripts in our transcriptome were tested for sequence similarity to the genomic and transcriptomic sequences of these species using local BLASTn. All transcripts with hits exceeding 70% identity at E-value below 10^-25^ were removed.

To assess the reliability of this procedure we searched the discarded sequences against known ciliate genomes (Extended Data Table 3) and did not obtain significant alignments for any sequence from 111 discarded OGGs. We further queried sequences from the remaining 1614 OGGs against the non-redundant Genbank database and did not obtain statistically significant alignments.

Finally, we checked for the presence of chimeric transcripts, i.e. transcripts assembled from reads originating from different organisms. We considered a transcript to be chimeric, if it yielded at least two BLAST hits (E-value < 10^-10^, Identity > 70%) that do not originate from the same organism with the overlap of at most 15 nt. No chimeric transcripts satisfying this criterion were identified.

### Phylogenetic analysis

To validate the phylogenetic tree obtained using 18S rRNA^28^ (Extended Data Figure 1), we also constructed a tree using concatenated coding sequences obtained in this study. The tree was built using paralog-free OGGs containing genes from all nine studied euplotid species with the sequences from *Tetrahymena thermophila* as an outgroup. The coding regions were aligned with MUSCLE^38^ and then the resulting alignments were concatenated. The tree was constructed with the Maximum Likelihood algorithm implemented in the PhyML package^53^ using the automatic model selection feature^54^ and 100 bootstrap replicates. Divergence times (per-branch *dS* values) were estimated with the m1 model of the PAML package. The ancestral sequences corresponding to frameshift sites were inferred using maximum parsimony (MP)^55,56^.

### Gains and losses of frameshift sites

To avoid misalignment errors we considered only frameshift sites within ±10 indel-free alignment blocks. The numbers of frameshift site gains and losses were calculated as the numbers of respective mutations, which all appear as insertions and deletions of thymines in stop codons (Figure 2a). Hence, the probability of frameshift site gain/loss at a specific context *i* was calculated as the number of gain/loss events *n* normalized over the number of the contexts *K*:

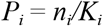

The contexts were defined by the codon preceding gained or lost stop codon, e.g. the context for frameshift sites is defined as NNN within NNN_TAR and in NNN_AR (potential gain upon insertion of T), R is purine (A or G), and the underscore separates codons in the coding reading phase.

### Calculating the inflation of frameshift sites

The temporal dynamics of the per-genome numbers of FSs is given by a logistic differential equation:

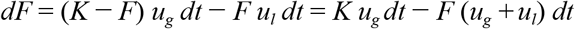

where *F*=*F*(*t*) is the (variable) number of FSs, *u_g_* and *u_l_* are the rates of site gain and loss, respectively, and *K* is the (constant) number of suitable ancestral contexts for the shift gain. The solution is:

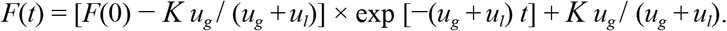

To simplify, let *A* = *Ku_g_* / (*u_g_*+ *u_l_*), *b* = *Ku_g_* and *F*(0) be the current number of frameshift sites at *t=0,* then:

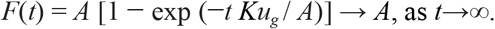

### Selection inference

We calculated the balance between selection and drift in the form of Kimura’s *S* defined as *S* = 4*sN_e_,* where *s* is the selection coefficient and *N_e_* is the effective population size.

Next, we derive the expression to infer *S* from the ratio of FS gain and loss probabilities. The observed mutation rate under mutation, selection, and drift is ^22^

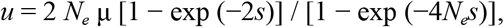

where μ is the neutral mutation rate.

The selection coefficient *s* can be presumed small, as events of frameshift site gain are observed. Thus, for *s*→0, 1 - exp (−2*s*) = 2*s*, and

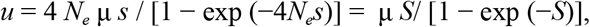

As we are dealing with rare evolutionary events, i.e., indels, and relatively small evolutionary times, the mutation rate may be presumed to equal the mutation probability: *u* = *P*.

Thus, the probability of FS gain is:

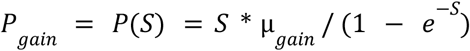

And, as the fitness effect of a frameshift site loss is equal to the negative fitness effect of a frameshift site gain, the probability of a frameshift site loss is:

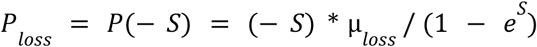

The analysis of thymine indels in our data (not shown) suggests that μ_*gain*_ = μ_*loss*_. Thus

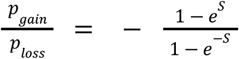

Or, alternatively:

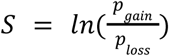

To obtain the lag load of the total body of frameshift sites, we first calculated the expected total number of FSs at equilibrium by normalizing the numbers of genes in our samples containing FSs to the total number of genes estimated for *E. focardii* and *E. octocarinatus* by Mozzicafreddo et al., 2021^51^ (Extended Data Figure 4). Next, by presuming independent fitness effects of each frameshift, we calculated the net lag load of frameshift sites as the product of fitness effects of all sites.

## Code availability

All scripts and data analysis protocols are available online at https://github.com/sofyagdk/euplotes.

## ACKNOWLEDGMENTS

PVB wishes to acknowledge financial support from SFI-HRB-Wellcome Trust Biomedical Research Partnership [Investigator in Science award 210692/Z/18/].

MAM was supported by the Russian Foundation for Basic Research, grant 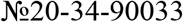.

JFA was personally supported by Irish Research Council Advanced Laureate grant IRCLA/2019/74.

AV thanks G. Di Giuseppe and P. Luporini for providing *Euplotes* species used in this study.

## EXTENDED DATA

**EXTENDED DATA FIGURE 1.**
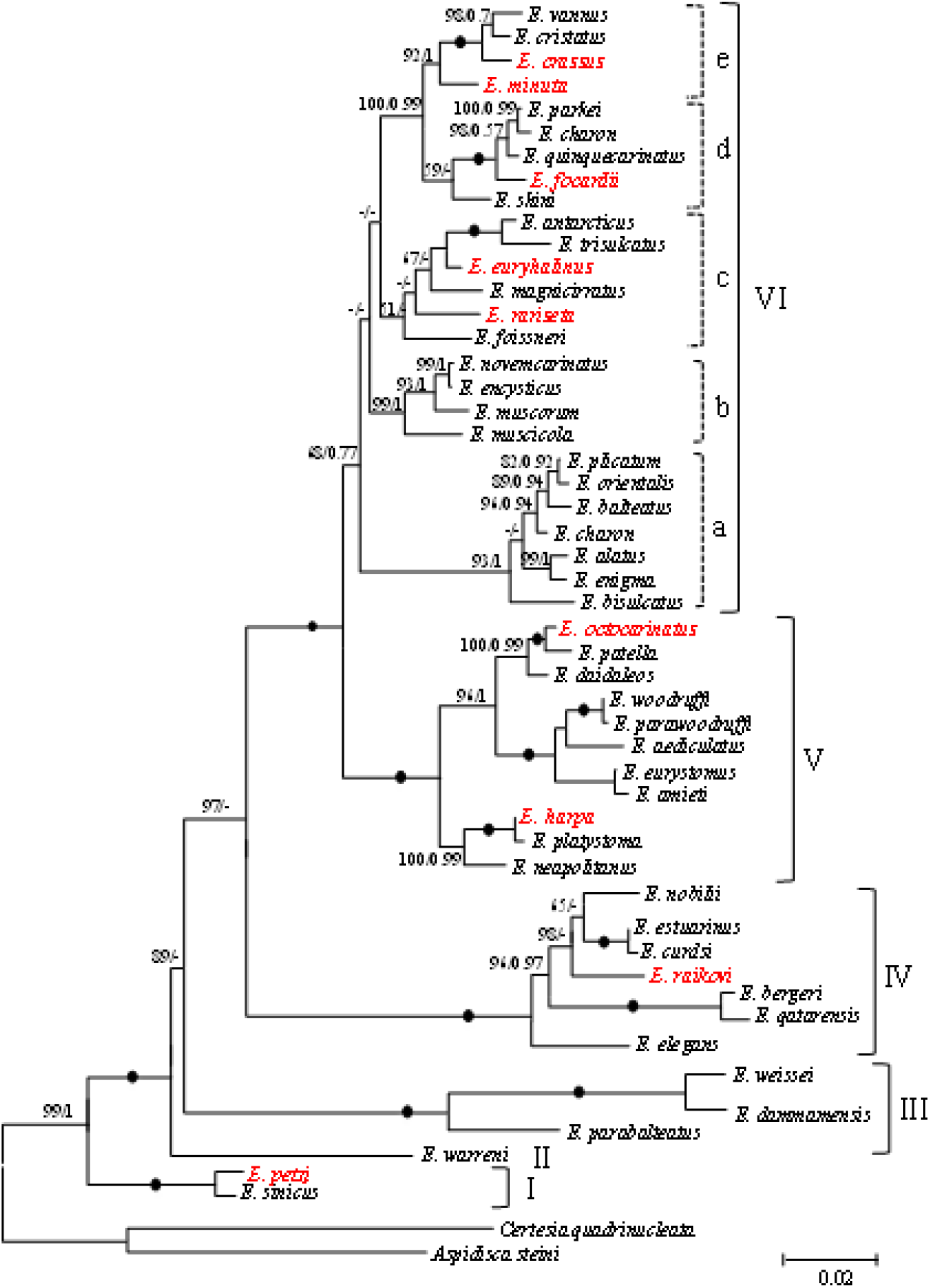
Phylogenetic tree based on 18S rRNA of different Euplotes. Euplotes species used in this work are shown in red. Numbers near the branches represent bootstrap values of Maximum Likelihood and Posterior Probability of Bayesian inference analyses, respectively. Fully supported (100/1.00) branches are marked with solid circles. All branches are drawn to scale. The scale bar corresponds to two substitutions per 100 nucleotide positions. Modified from Valbonesi et al., 2021^26^

**EXTENDED DATA FIGURE 2.**
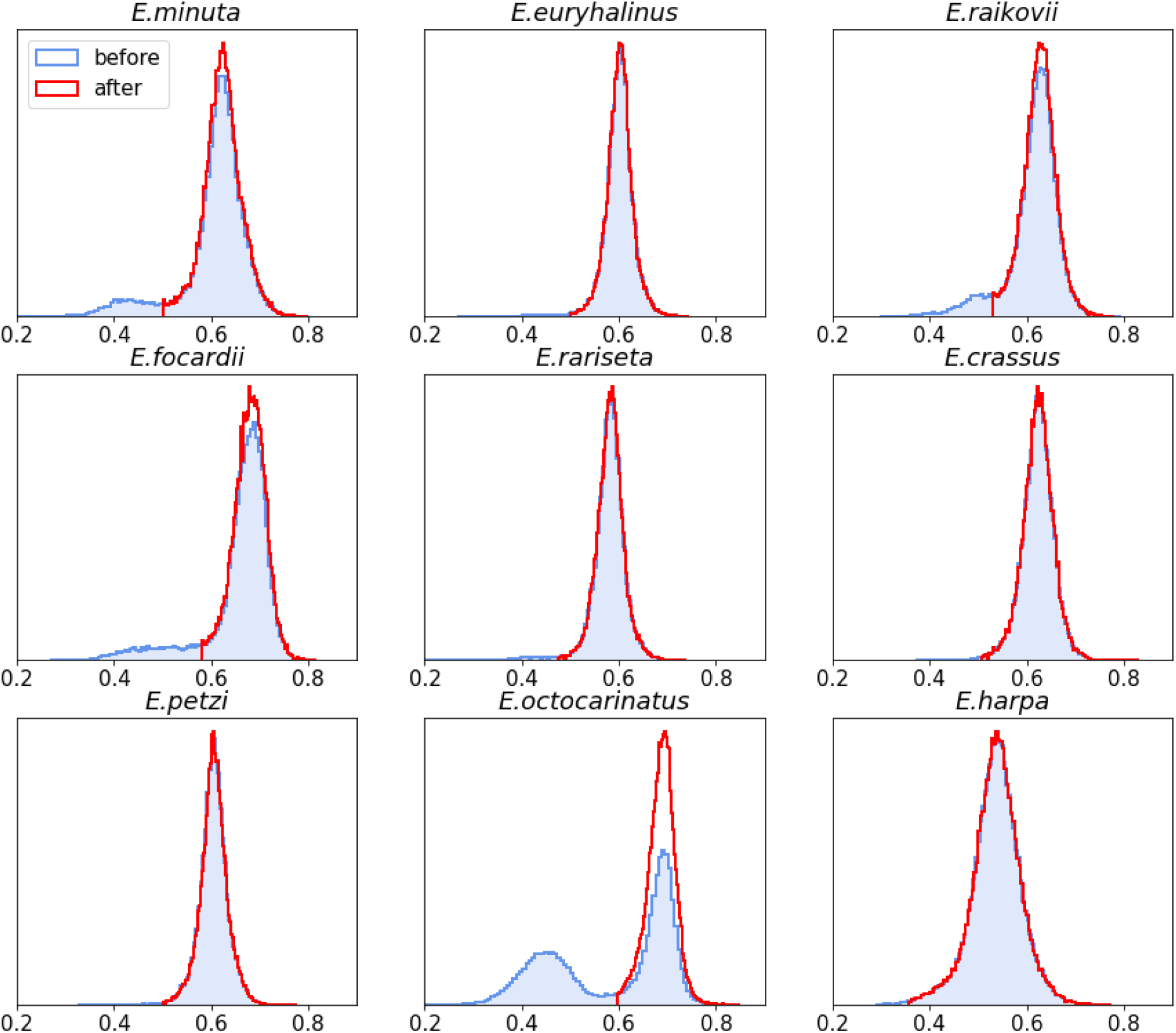
Transcriptome quality assessment based on AT content. Distributions of AT-compositions calculated in each transcript for each *Euplotes* species before (blue) and after filtration (red), see Methods.

**EXTENDED DATA FIGURE 3.**
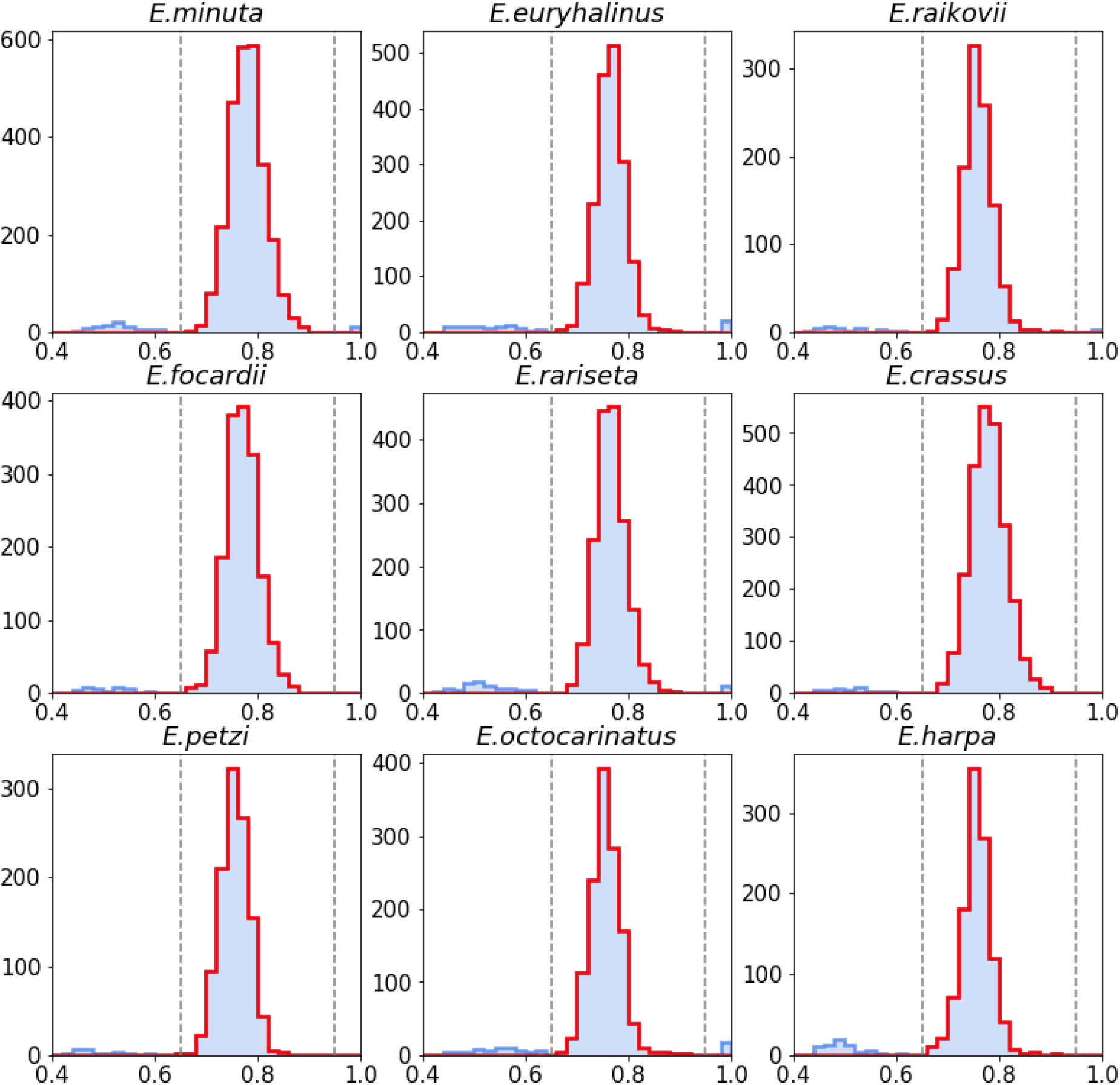
Transcriptome quality assessment based on sequence similarity with orthologs. Distributions of pairwise nucleotide identity for transcripts before (blue) and after (red) filtration for each *Euplotes* species.

**EXTENDED DATA FIGURE 4.**
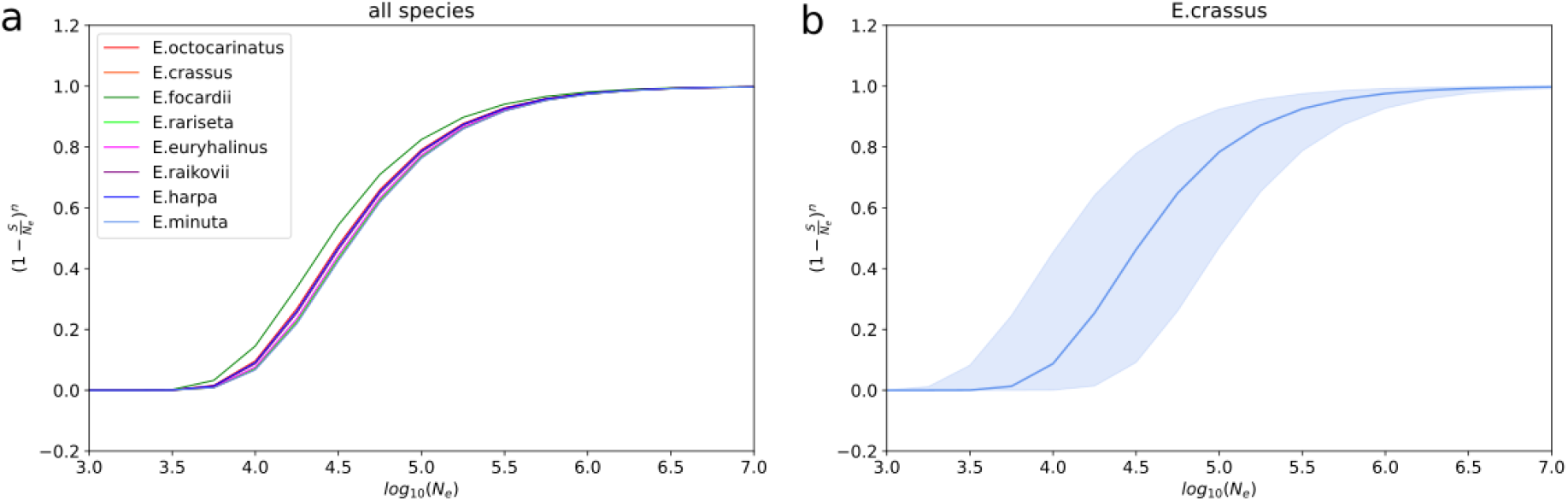
Selection against the total body of euplotid frameshift sites when the equilibrium numbers of frameshifts are reached. **(a)** Dependency between effective population size and normalized lag loads conveyed by frameshifts in all considered *Euplotes* species **(b)** Confidence interval for the dependency shown on panel (a) for *E. crassus*.

**EXTENDED DATA FIGURE 5.**
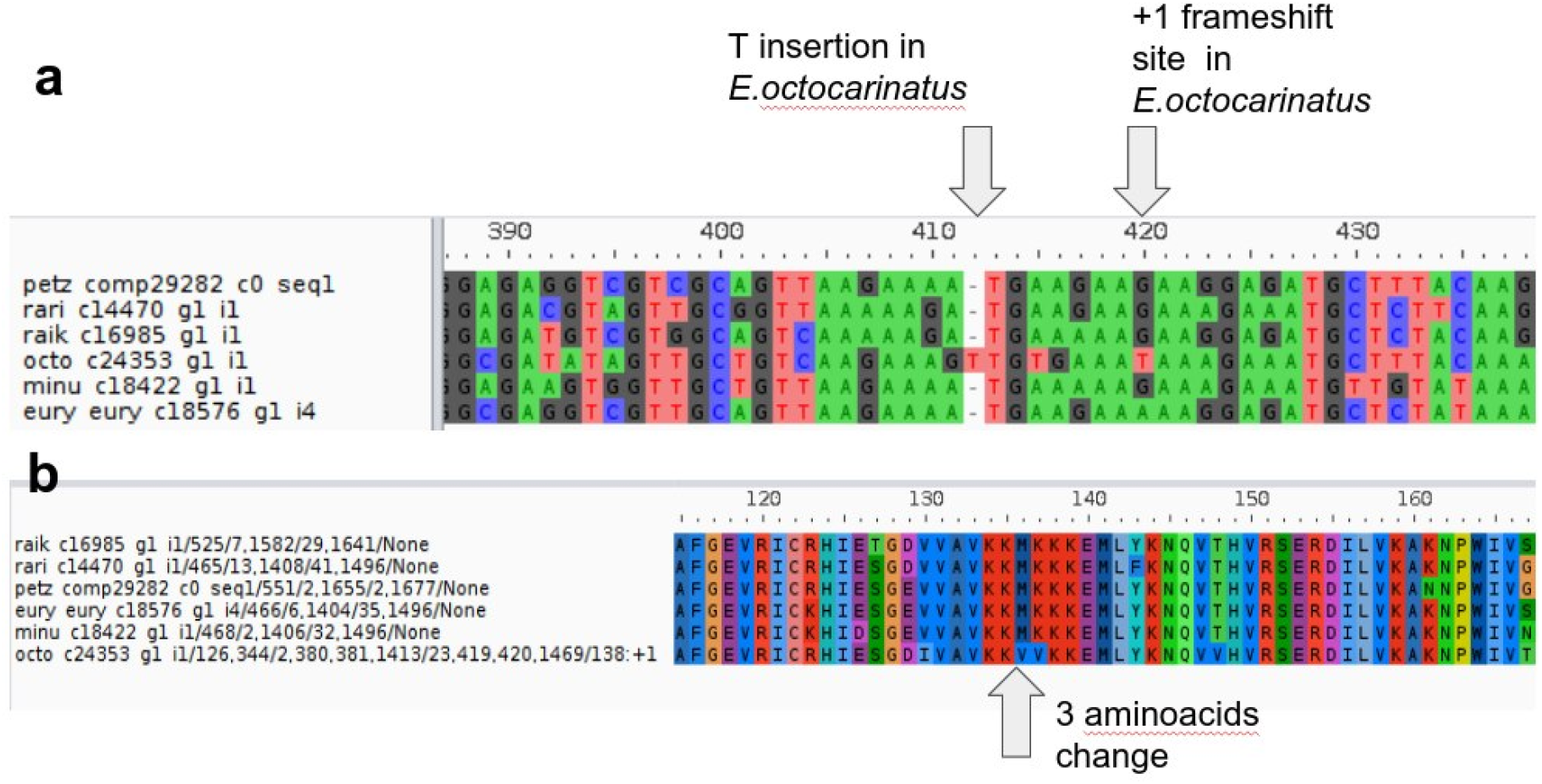
An example of restoration of the initial reading frame through ribosomal frameshift. **(a)** Single-nucleotide insertion in *E. octocarinatus* and a +1 frameshift site at position 420 restoring the initial reading frame **(b)** Respective protein sequence alignment. Note the substitution MK dipeptide with VV on coordinates 135-136.

